# Analysis of Genetically Regulated Gene Expression identifies a trauma type specific PTSD gene, SNRNP35

**DOI:** 10.1101/581124

**Authors:** Laura M Huckins, Michael S Breen, Chris Chatzinakos, Jakob Hartmann, Torsten Klengel, Ana C da Silva Almeida, Amanda Dobbyn, Kiran Girdhar, Gabriel E Hoffman, Claudia Klengel, Mark W Logue, Adriana Lori, Filomene G Morrison, Hoang T Nguyen, Yongjin Park, Douglas Ruderfer, Laura G Sloofman, Sanne JH van Rooij, PTSD Working Group of Psychiatric Genomics Consortium, Dewleen G Baker, Chia-Yen Chen, Nancy Cox, Laramie E Duncan, Mark A Geyer, Stephen J. Glatt, Hae Kyung Im, Adam X Maihofer, Victoria B Risbrough, Jordan W Smoller, Dan J Stein, Rachel Yehuda, Israel Liberzon, Karestan C Koenen, Tanja Jovanovic, Manolis Kellis, Mark W Miller, Silviu-Alin Bacanu, Caroline M Nievergelt, Joseph D Buxbaum, Pamela Sklar, Kerry J Ressler, Eli A Stahl, Nikolaos P Daskalakis

## Abstract

PTSD has significant genetic heritability; however, it is unclear how genetic risk influences tissue-specific gene expression. We used brain and non-brain transcriptomic imputation models to impute genetically regulated gene expression (GReX) in 9,087 PTSD-cases and 23,811 controls and identified thirteen significant GReX-PTSD associations. The results suggest substantial genetic heterogeneity between civilian and military PTSD cohorts. The top study-wide significant PTSD-association was with predicted downregulation of the Small Nuclear Ribonucleoprotein U11/U12 Subunit 35 (SNRNP35) in the BA9 region of the prefrontal cortex (PFC) in military cohorts. In peripheral leukocytes from 175 U.S. Marines, the observed PTSD differential gene expression correlated with the predicted blood GReX differences for these individuals, and deployment stress downregulated *SNRNP35* expression, primarily in Marines with post-deployment PTSD. SNRNP35 is a subunit of the minor spliceosome complex and *SNRNP35* knockdown in cells validated its functional importance in U12-intron splicing. Finally, mimicking acute activation of the endogenous stress axis in mice downregulated PFC *Snrnp35* expression.

## INTRODUCTION

Trauma exposure is ubiquitous, particularly in veterans and impoverished, high-risk civilian populations. Post-traumatic stress disorder (PTSD) is a debilitating psychiatric condition, occurring in some individuals exposed to trauma, while the large proportion of individuals do not experience PTSD, and remain resilient even after repeated, prolonged or severe exposure to trauma (Bonanno, 2004; Kessler et al., 2005; Kessler et al., 1995). Understanding which individuals may be susceptible or resilient to PTSD is vital in the development of effective interventions and treatments. Twin studies have repeatedly demonstrated that PTSD diagnosis and symptoms are heritable, with heritability estimates ranging from 30-71% (Daskalakis et al., 2018b; Nievergelt et al., 2018a), in line with other psychiatric disorders. Several genome-wide association studies (GWAS) have identified genetic variants or loci associated with PTSD susceptibility, although most of the associations have failed to replicate (Daskalakis et al., 2018b; Nievergelt et al., 2018a). Most recently, a GWAS by the Psychiatric Genomics Consortium for PTSD (PGC-PTSD) demonstrated SNP-based heritability of 21%, comparable to other psychiatric disorders, and demonstrated genetic correlations with schizophrenia, bipolar disorder, and major depressive disorder (Duncan et al., 2018).

Despite the substantial success of GWAS in elucidating the genetic etiology of psychiatric disorders, resulting associations may be difficult to interpret biologically. At best, these studies result in large lists of associated loci, which require careful curation to prioritize genes (Visscher et al., 2017). On the other hand, studies of the transcriptome may yield more readily biologically interpretable results. However, progress in these studies in hampered by small sample sizes, due in part to the cost and inaccessibility of the primary tissue of interest, i.e. brain. Transcriptomic Imputation (TI) approaches leverage large reference transcriptome data sets, e.g. the Genotype-Tissue Expression (GTEx) project and CommonMind Consortium (CMC), in order to estimate relationships between genotypes and gene expression, and to create predictor models of genetically regulated gene expression (GReX) (Gamazon et al., 2015; Gusev et al., 2016). TI algorithms, thus, allow us to identify genes with predicted disease-associated differential GReX in specific tissue. These analyses allow us to probe gene expression in vastly larger sample sizes, yielding sufficient power to detect genes with small effect sizes (Gamazon et al., 2015), which represent a substantial proportion of the risk for complex diseases (Fromer et al., 2016).

Notably, PTSD development and symptom trajectories differ according to trauma type. For example, the prevalence of PTSD differs significantly between rape survivors (45%), combat veterans (30%) and following natural disasters (4%) (Kessler et al., 2017; Kessler et al., 2005; Yehuda et al., 2015). Both in civilian and military populations, index trauma type and exposure severity significantly predict PTSD diagnosis, symptom severity, and severity of severity of specific symptom clusters (Graham et al., 2017; Jakob et al., 2017; Kessler et al., 2005; Prescott, 2012). While the differential prevalence, symptoms, and outcomes have been characterized in depth, to our knowledge no study has investigated differing genetic underpinnings of PTSD according to trauma type. The current PGC-PTSD study includes large collections of both military PTSD (M-PTSD) and civilian PTSD (C-PTSD) cohorts. Although these are only a proxy for trauma type (e.g., combat veterans can experience non-combat military trauma and trauma in their civilian life, and both military and civilian trauma cohorts include a wide range of trauma types) and a number of individual factors in addition to trauma type may differentiate these cohorts, they provide a powerful opportunity to begin to probe differing genetic etiologies of PTSD between these two groups.

## RESULTS

### Genetically regulated gene expression in brain and non-brain tissues is associated with PTSD

We imputed GReX across 32,898 individuals (9,087 cases and 23,811 controls; demographics in **Table S1A**; analytic plan in Figure S1) from the largest multi-cohort PTSD GWAS, collected by PGC-PTSD and tested for association with case-control status. Since PTSD development involves multi-systemic dysregulation (Daskalakis et al., 2018a; Sareen, 2014), we imputed GReX in 22 tissues, using GTEx- and CMC- derived predictor models (11 brain regions, five cardiovascular tissues, three endocrine tissues, one peripheral nerve, subcutaneous adipose tissue and whole blood).

No genes reached study-wide significance (p=2.36×10^−7^; based on 211,466 tissue-gene pairs, across all tissues) in our overall trans-ethnic meta-analysis (Manhattan plot in Figure 1A; association statistics and gene names in Table 1), including African American (AA), European American (EA), Latino or Hispanic American (LA) and South African (SA) cohorts as in (Duncan et al., 2018). However, *DLG4* (Pituitary, p=4.51×10^−7^), encoding for postsynaptic density protein 95 (PSD-95), and *SNRNP35* (Prefrontal cortex (PFC) BA9, p=4.25×10^−6^), encoding for Small Nuclear Ribonucleoprotein U11/U12 Subunit 35, reached within-tissue significance (5.24×10^−6^ based on an average of 9542 genes in each tissue; exact statistical thresholds in Table S2). In our EA-specific meta-analysis (5,236 cases and 13,357 controls), *SNRNP35* reached study-wide significance (PFC BA9, p=5.47×10^−08^).

**Table 1.** Study-wide and tissue-wide significant GReX associations with PTSD.

| # | Trauma type (proxy) | Females (%) | Ancestry | TOTAL (cases/controls) | Tissue | Gene Symbol | Gene Name | Direction, Z-score | P |
| --- | --- | --- | --- | --- | --- | --- | --- | --- | --- |
| 1 | ALL | 30.3% | ALL | 32,898 (9,087/23,811) | Pituitary | *DLG4* | discs large MAGUK scaffold protein 4 (also known as postsynaptic density protein 95) | ↑, 5.05 | 4.51x10<sup>-7</sup> |
| 2 | ALL | 30.3% | ALL | 32,898 (9,087/23,811) | Frontal Cortex BA9 | *SNRNP35* | small nuclear ribonucleoprotein U11/U12 subunit 35 | ↓, -4.60 | 4.25x10<sup>-6</sup> |
| 3 | ALL | 20.0% | EUR | 18,593 (5,236/13,357) | Frontal Cortex BA9 | ***SNRNP35\**** | small nuclear ribonucleoprotein U11/U12 subunit 35 | ↓, -5.44 | 5.47 x10<sup>-08 a</sup> |
| 4 | MILITARY | 10.7% | ALL | 19,959 (6,109/13,850) | Frontal Cortex BA9 | ***SNRNP35\**** | small nuclear ribonucleoprotein U11/U12 subunit 35 | ↓, -5.35 | 8.85 x10<sup>-08</sup> |
| 5 | MILITARY | 10.7% | ALL | 19,959 (6,109/13,850) | Heart Atrial Appendage | ***SENPI\**** | SUMO1/sentrin specific peptidase 1 | ↑, 5.19 | 2.08 x10<sup>-07 b</sup> |
| 6 | MILITARY | 10.7% | ALL | 19,959 (6,109/13,850) | Anterior Cingulate Cortex BA24 | *NEDD9* | neural precursor cell expressed, developmentally down-regulated 9 | ↑, 4.66 | 3.18 x10<sup>-06</sup> |
| 7 | MILITARY | 9.2%<sup>#</sup> | EUR | 13,281 (4,099/9,182) | Frontal Cortex BA9 | ***SNRNP35\**** | small nuclear ribonucleoprotein U11/U12 subunit 35 | ↓, -6.08 | 1.23 x10<sup>-09 c</sup> |
| 8 | MILITARY | 9.2%<sup>#</sup> | EUR | 13,281 (4,099/9,182) | Heart Atrial Appendage | *SENPI* | SUMO1/sentrin specific peptidase 1 | ↑, 4.76 | 1.92 x10<sup>-06 d</sup> |
| 9 | MILITARY | 9.2%<sup>#</sup> | EUR | 13,281 (4,099/9,182) | Cerebellar Hemisphere | *RPS3* | ribosomal protein S3 | ↓, -4.69 | 2.77 x10<sup>-06</sup> |
| 10 | MILITARY | 18.0%<sup>#</sup> | AA | 3,327 (1,204/2,123) | Hypothalamus | *SYNGR2* | synaptogyrin 2 | ↑, 4.69 | 2.61 x10<sup>-06</sup> |
| 11 | MILITARY | 18.0%<sup>#</sup> | AA | 3,327 (1,204/2,123) | Nucleus Accumbens | *CNOT1* | CCR4-NOT transcription complex subunit 1 | ↑, 4.66 | 3.25 x10<sup>-06</sup> |
| 12 | CIVILIAN | 54.8% | EUR | 5,312 (1,137/4,175) | Nucleus Accumbens | *GCM1* | glial cells missing homolog 1 | ↑, 4.81 | 1.53 x10<sup>-06</sup> |
| 13 | CIVILIAN | 60.0% | AA | 7,243 (1,711/5,532) | Cortex | *MARCH11* | membrane associated ring-CH-type finger 11 | ↓, -4.64 | 3.37 x10<sup>-06</sup> |

Notes. All genes have within-tissue significance. Raw data were not available for military cohorts; for these, sample proportions are extrapolated from previous publications of these data. Z-score corresponds to the effect of differences in genetically regulated gene expression (GReX) in cases vs. controls. Standard errors are not scaled to the same unit variance between military and civilians; as such, military and civilian analyses were combined using sample-size based meta-analysis. See **Table S1** for sample collection details. Bold lettering with \* denotes genes with study-wide significance. <sup>a</sup> male-specific meta-analysis: N cases= 3512, N controls= 11769, z = -5.48; P=4.20 x10<sup>-08</sup>. <sup>b</sup> male-specific meta-analysis: N cases= 4578, N controls=14770, z = 4.67; P=2.94 x10<sup>-06</sup>. <sup>c</sup> male-specific meta-analysis: N cases= 3367, N controls=10754, z = -5.67; P=1.41 x10<sup>-08</sup>. <sup>d</sup> male-specific meta-analysis: N cases= 3367, N controls=10754, z = 4.64; P=3.39 x10<sup>-06</sup>.

**Figure 1.**
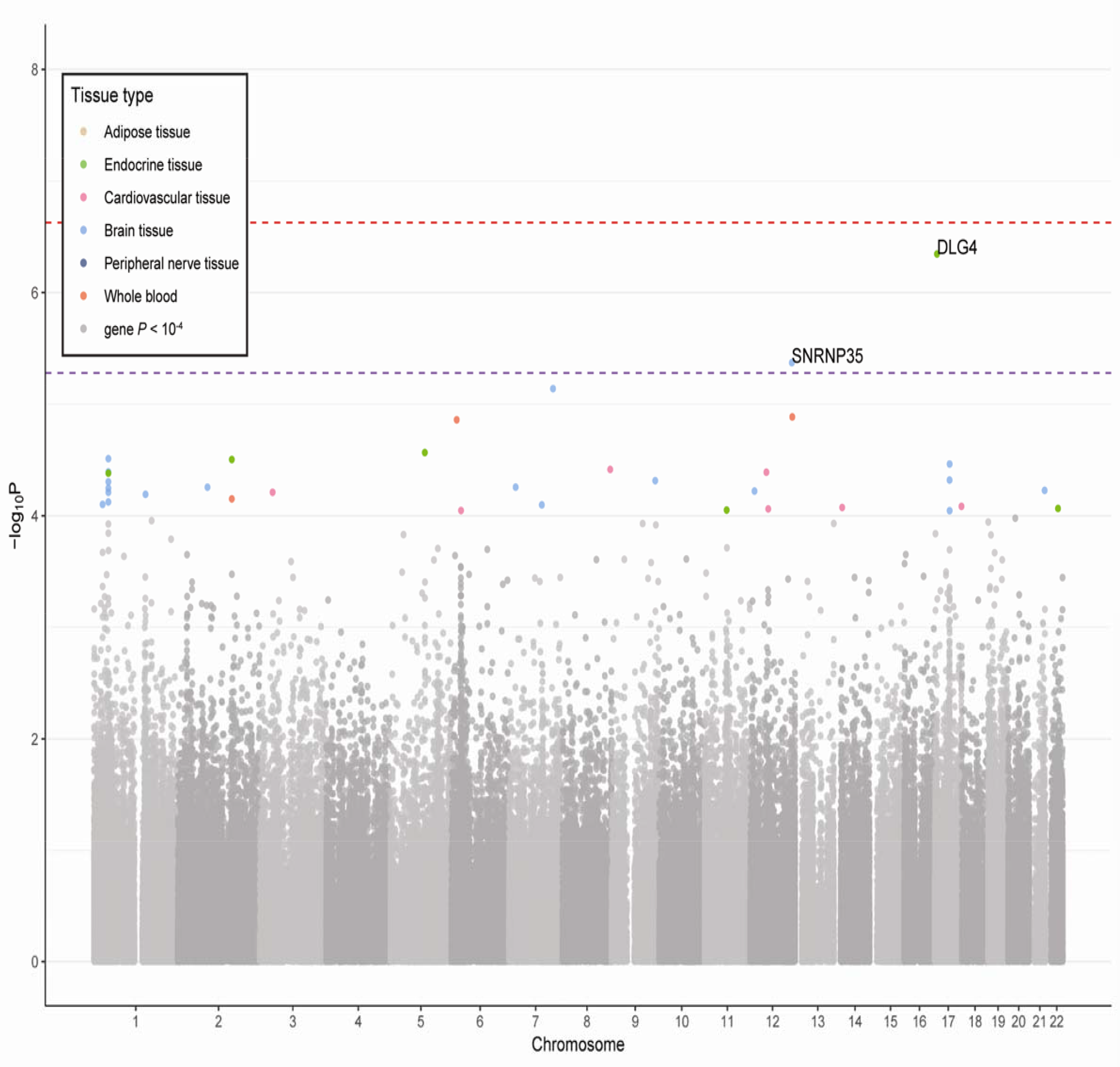
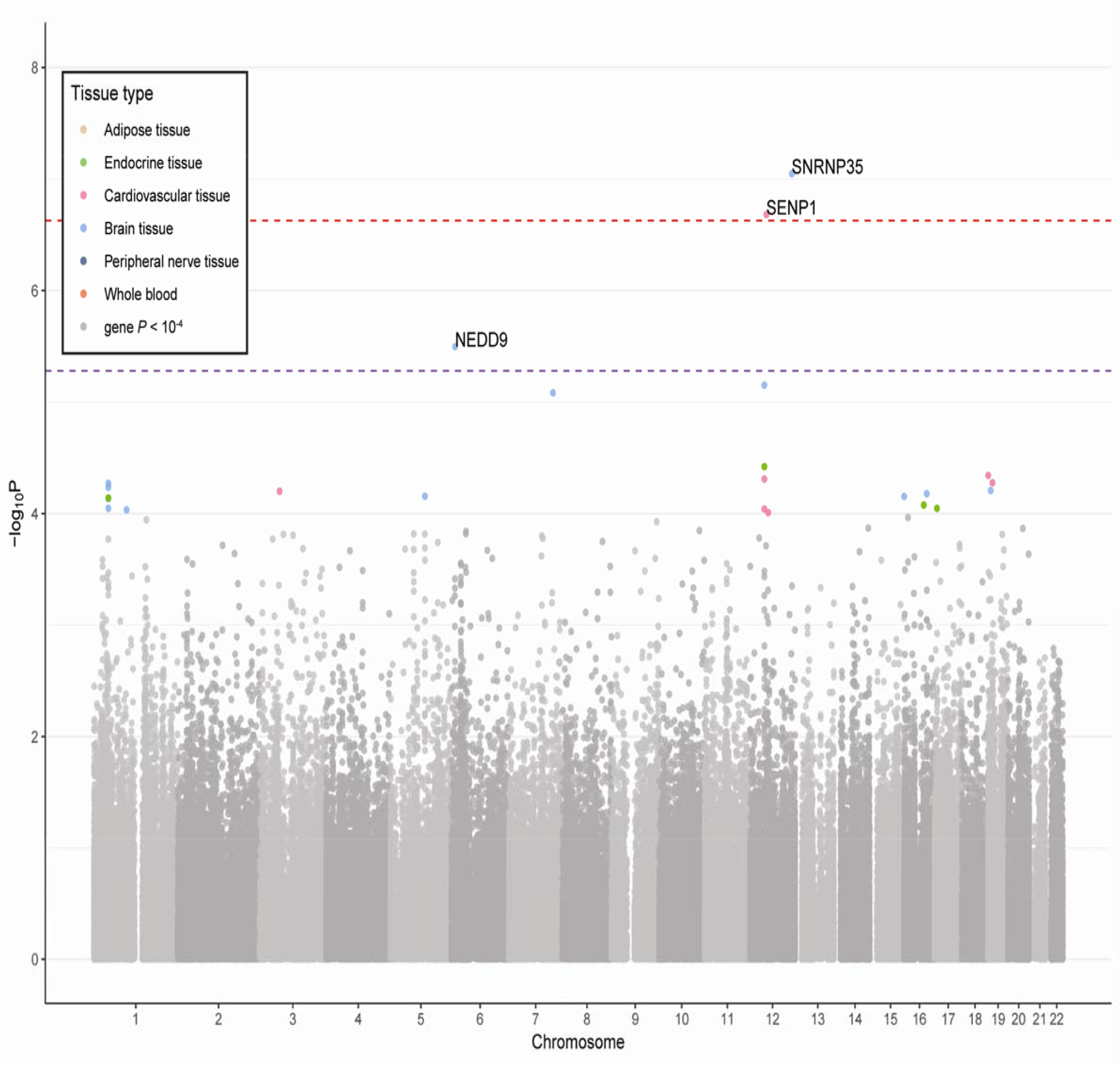
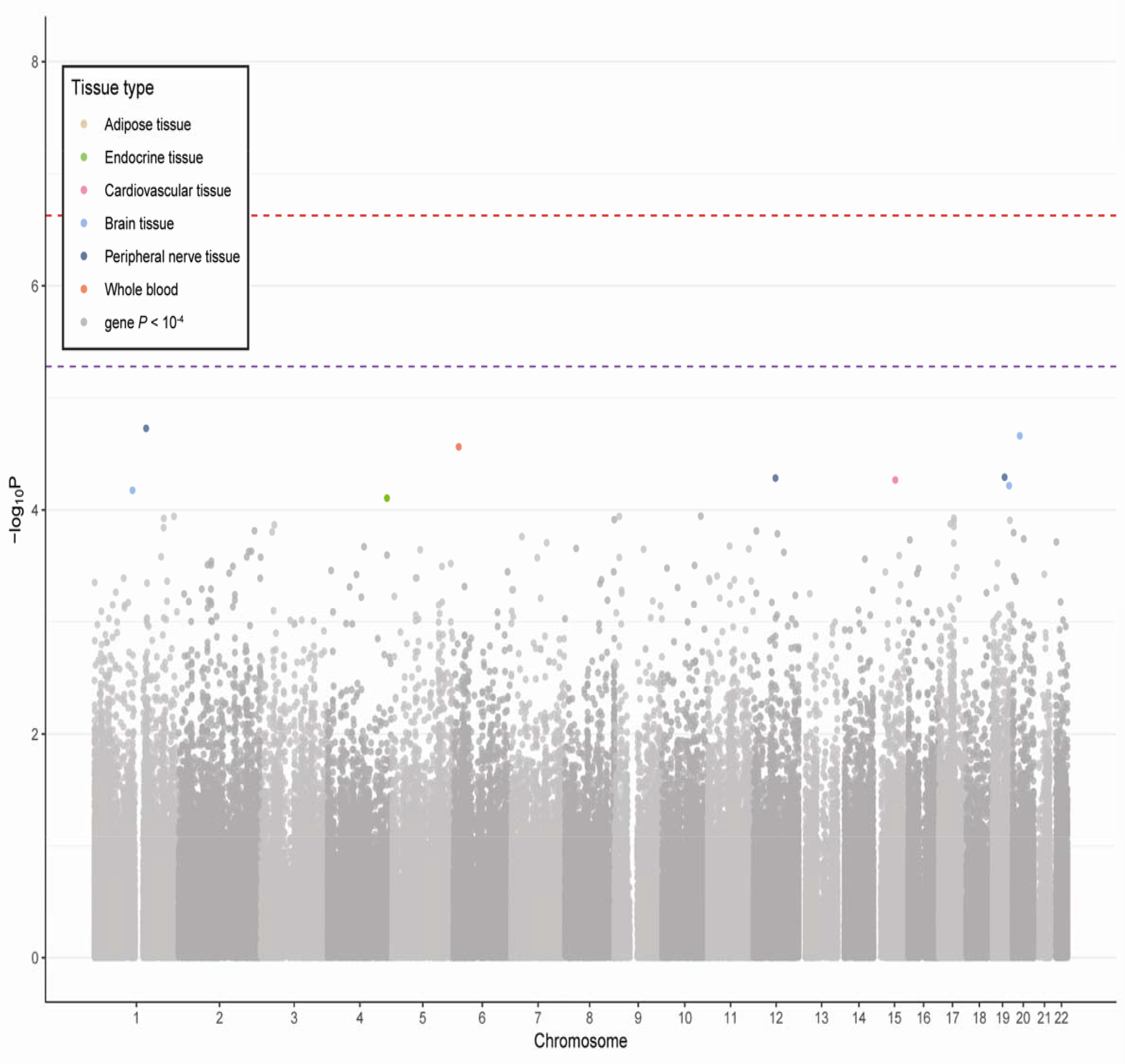
Gene-by-tissue associations with PTSD. (A) No associations reach study-wide significance (depicted by red discontinuous line) in our overall transethnic meta-analysis, but two genes reach within-tissue significance (depicted by purple discontinuous line). (B) Two genes reach study-wide significance in the military-only meta-analysis and one reached within tissue significance; (C) while no genes are significant at any of the significance thresholds in the civilian-only meta-analysis. Gene-tissue pairs are color-coded according to tissue type.

### GReX associations differ between Military and Civilian PTSD cohorts

We next hypothesized that the genetic architecture of PTSD may differ according to trauma type. The genetic signatures underlying predisposition to PTSD following combat or other military-specific trauma may be distinct from those underlying PTSD following motor vehicle accident, neighborhood or domestic violence and sexual assault. We therefore stratified our meta-analysis according to trauma type, using military and civilian cohorts as proxies. Our M-PTSD meta-analysis, comprised of 6,109 cases and 13,850 controls, identified two genes reaching study-wide significance; *SNRNP35* (PFC BA9, p=8.85×10^−8^) and *SENP1* (Atrial Appendage, p=2.08×10^−7^), both on chromosome 12 (Figure 1B). A third gene reached within-tissue significance; *NEDD9* (Anterior Cingulate Cortex BA24, p=3.18×10^−6^). When restricting our analysis to only military cohorts of EA descent, the *SNRNP35* association remained study-wide significant, while the *SENP1* association reached only within-tissue significance in the same analysis (Table 1). A third gene, *RPS3*, reached within-tissue significance in this EA analysis, as did two further genes, *SYNGR2* and *CNOT1*, in the AA analysis.

We did not identify any genes reaching study-wide significance in our C-PTSD transethnic analysis (2,978 cases and 9,961 controls; Figure 1C), which may be, in part, due to its lower power and smaller sample size compared to the respective military analysis. Two genes (*GCM1* and *MARCH11)* reached within-tissue significance in EA- and AA-specific C-PTSD analysis, respectively (Table 1). Notably, the genes reaching study-wide significance in our military analysis do not approach significance in our civilian analysis; rather, as demonstrated in Figure1B-C, these associations seem to be trauma type specific.

### Trauma-type specific associations are not driven by gender differences

It is possible that these proxy trauma types, and the resulting different association patterns, are confounded by sex. Notably, the M-PTSD sample is primarily male (~90%). We hypothesized that, if these association patterns are indeed driven by sex, we should see an enrichment of shared nominally-significant associations and greater correlation of association statistics within-sex compared to within trauma type.

To test this hypothesis, we compared four analyses: male C-PTSD, female C-PTSD, male M-PTSD, and female M-PTSD, for all 22 tissues. We saw significant enrichment of nominally significant male C-PTSD associations in female C-PTSD, and vice-versa (binomial tests *P*-value < 5.9×10^−8^), but not in any of the other pairwise enrichments (**Table S3A**). Further, (**Table S3B**), male C-PTSD and male M-PTSD association statistics were uncorrelated (average across-tissue Pearson correlation and *P*-value: r=0.001, p=0.457), while male C-PTSD and female C-PTSD association statistics had a significant relationship (r=0.138, p=4.4×10^−42^). Despite this broad lack of support for the hypothesis of sex differences driving the trauma type specific signals, we note that our top M-PTSD association, *SNRNP35*, reached study-wide significance in male-specific meta-analyses: of EA cohorts (Table S4; 3512 cases/ 11769 controls) and of EA M-PTSD cohorts (3367 cases/ 10754 controls), but not in any of our female analyses, implying that this association may have trauma type, ancestry and sex specificity. Our male-specific meta-analyses also identified additional three genes (including *SENP1*) reaching tissue-specific significance, and the female-specific analyses identified two other genes (Table S4). However, none of these genes were significant in both the male and female meta-analyses, and, given the small sample sizes of some of our meta-analyses, especially male C-PTSD and female M-PTSD (**Table S1B**), we urge that sex-specific analyses be pursued further in larger datasets.

### GReX associations with PTSD are enriched for known and novel pathways

We used MAGMA (de Leeuw et al., 2015) to test for gene-set enrichment in our overall, C-PTSD and M-PTSD transethnic meta-analyses’ results. For each analysis, we tested for enrichment of a set of PTSD candidate genes identified in previous PTSD literature (**Table S5)**, 92 hypothesis-driven gene-sets and ~8,500 publicly available gene-sets (**Supplementary Materials**). First, our literature-derived PTSD candidate genes were significantly enriched in the C-PTSD analysis (Table S6; p=0.0014), but not the overall, or M-PTSD analyses. The top individual genes included *RIN1* (p=1.94×10^−4^), *GJA1* (p=2.52×10^−4^), *BDNF* (p=3.90×10^−4^), *OXTR* (p=4.24×10^−4^) and *DNMT1* (p=8.71×10^−4^); none with within-tissue significance.

We identified two gene-sets that were significantly enriched in the overall analysis (Table S6): protein phosphatase type 2A regulator activity (p=2.26×10^−6^) and uric acid levels (p=9.94×10^−5^). We then identified 11 significantly enriched gene-sets (with FDR<0.1) in the C-PTSD analysis (Table S6), including, from our hypothesis-driven analysis: genes intolerant to loss-of-function mutations (p=9.98×10^−5^), PSD-95 gene-set (p=0.0053), genes with loss-of-function mutations in autism (p=0.0058) and two circadian rhythm pathways (p<0.0063). Our agnostic analysis identified further gene-sets related to head shortening and head dysmorphology in mice (p<1.98×10^−5^), and early phase of HIV Life Cycle (p=2.22 ×10^−5^).

Finally, we identified 29 gene-sets significantly enriched in the M-PTSD analysis (Table S6). Only one gene-set from the hypothesis-driven analysis was significant; H3K27 acetylation peaks specific to dorsolateral PFC (DLPFC) neurons (p= 6.4×10^−4^). 28 gene-sets from our agnostic analysis were significant; these included astrocyte differentiation (p= 1.51×10^−7^), two olfactory pathways (p< 5.55×10^−7^), multiple pathways associated with glia functions (p< 5.14×10^−5^), decreased motor neuron number (p= 1.94×10^−5^), two pathways related to protein tyrosine kinase activity (p< 5.15×10^−5^) and two pathways related to RNA stability (p< 1.15×10^−4^).

### Predicted PTSD GReX differences are concordant with the observed PTSD gene expression differences

We sought to validate our GReX results in a subset of our PGC-PTSD samples for which observed peripheral leukocyte gene expression data was generated as part of a large prospective U.S. Marine cohort, the Marine Resiliency Study (MRS; **Supplementary Materials**, Figure S2). Clinical interviews and peripheral blood samples were collected from U.S. Marines one month prior-to deployment (*i.e.* pre-deployment, N=175) and three-months following exposure to conflict zones (*i.e.* post-deployment, N=157). We performed, at both time points, a differential expression analysis based on post-deployment PTSD, covarying for genetic ancestry, age, traumatic brain injury (TBI), alcohol and nicotine. We identified 280 genes nominally associated with future development of PTSD at pre-deployment, and 160 genes at post-deployment (*p* <0.05; **Table S7**).

In parallel, we carried out differential expression analysis on all GReX imputed tissues, and on all paired samples for which observed blood gene expression was available using a matching strategy (**Tables S8A-B)**. We then examined the association between directionality of change statistics (log fold-change) for the observed differential expression and predicted differential GReX. The correlations between observed and whole blood GReX fold-changes were significant at both pre- (*r*=0.45, *p*=1.39×10^−7^, Figure 2A, **Table S9**) and post-deployment (*r*=0.47, *p*=3.09×10^−8^, Figure 2B, **Table S9**). We also found the strongest concordance of observed PTSD effects with predicted effects on whole-blood GReX, compared to GReX of all other tissues, validating not only the accuracy and tissue-specificity of our TI-based approach (Figure 2C-D; **Table S9**).

**Figure 2.**
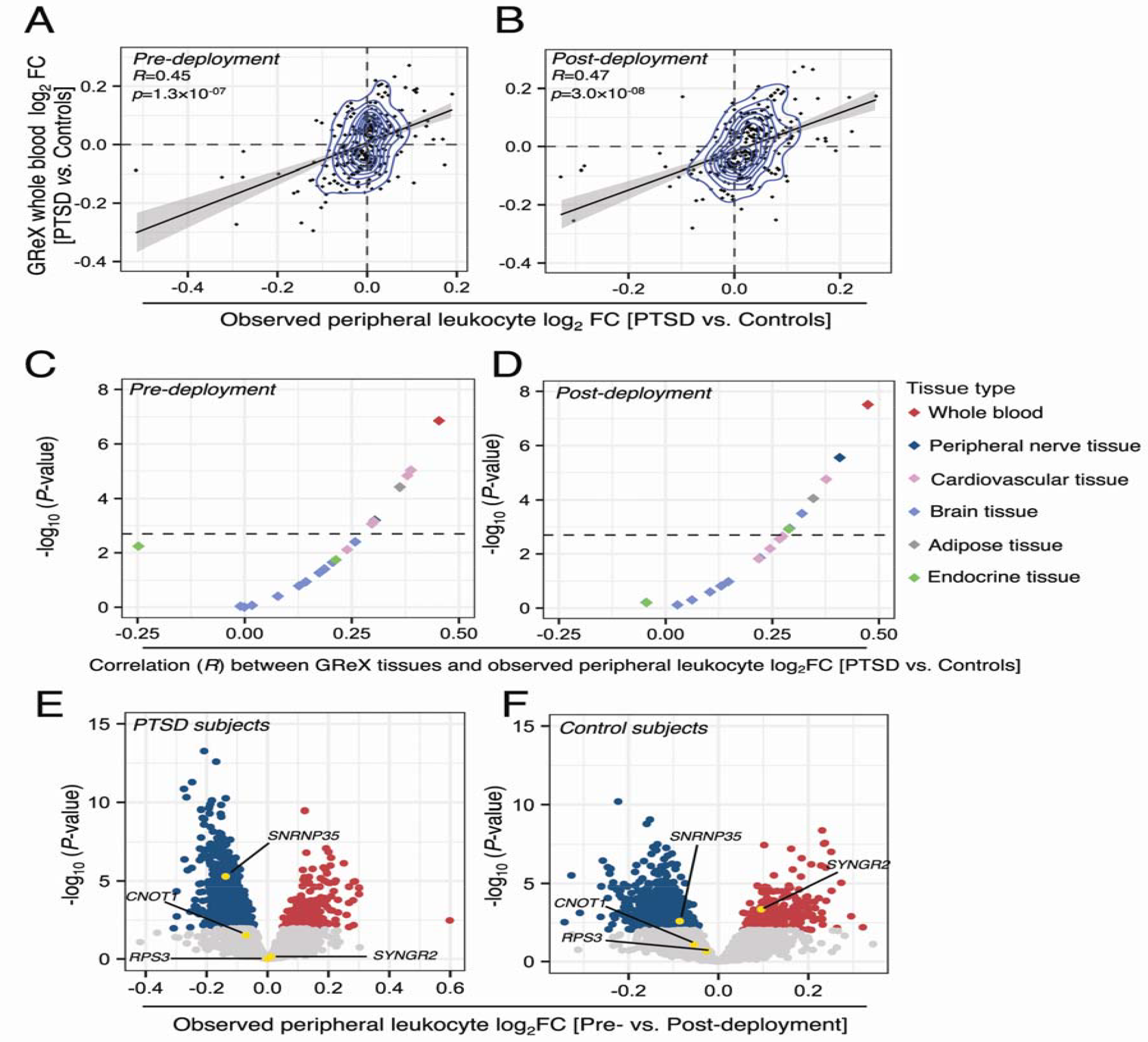
PTSD genetically-regulated expression (GReX) differences are concordant with the observed blood PTSD gene expression differences in the Marine Resiliency Study. Correlation between PTSD (vs. controls) fold changes (FC) of observed peripheral leukocyte gene expression in x-axis, measured at pre-(A) and post-deployment (B), and whole blood genetically-regulated expression (GReX) in y-axis. Correspondence between the coefficients of correlation between peripheral leukocyte gene expression and GReX across multiple tissues (x-axis) and level of correspondent significance (y-axis) at pre-(C) and post-deployment (D). Volcano-plot of the post-vs. pre-deployment differential expression changes in subjects with (E) and without (F) post-deployment PTSD. Points in panels C and D are color-coded according to tissue type. Red dots in panel E and F depict significantly upregulated at FDR significance threshold, while the blue depict significantly downregulated at FDR significance threshold.

### *SNRNP35* is downregulated in US Marines following deployment

The longitudinal design of the MRS enabled us to compare baseline to post-deployment peripheral leukocyte gene expression, separately for PTSD cases and control samples. We identified 1335 genes with FDR-significant longitudinal changes in expression in PTSD cases and 1161 genes in control samples (**Table S10**). *SNRNP35*, which was the gene with the most significant GReX associations with M-PTSD (Table 1), was found to be downregulated in response to deployment stress more strongly in PTSD cases than in control samples (FC= −0.137, *p*=0.0025 - Figure 2E; FC= −0.086, *p*=0.02 - Figure 2F, respectively), concordant with the direction of effect in our PrediXcan analysis.

### *SNRNP35* is part of RNA-processing gene networks in DLPFC

Our results emphasized the potential role of *SNRNP35* gene expression in BA9 brain region, which together with BA46 comprise DLPFC. Co-expression network analysis of a large CMC DLPFC RNA-seq dataset (Fromer et al., 2016) of healthy subjects (N=279) revealed that SNRNP35 is part of a co-expression sub-network/module, containing 152 genes (https://www.synapse.org/#!Synapse:syn7118802). This module is enriched with a range of RNA binding and processing functions (GO biological processes, molecular functions and cellular components), highlighting module functional specificity related to RNA binding (adjusted *P*-value =2.78×10^−16^) and RNA processing/splicing (adjusted *P*-value =2.01×10^−08^; **Table S11**-**12**).

### SNRNP35 knockdown reduces U12 splicing

SNRNP35 protein is subunit of the minor spliceosome, which catalyzes the removal/splicing of an atypical class of introns – U12-type (0.5% of all introns), from messenger RNAs (mRNAs) (Turunen et al., 2013). We tested whether *SNRNP35* downregulation is sufficient to cause a functional impact on U12 splicing in cell-culture experiments. Using small hairpin RNAs (shRNAs), we specifically tested the effect of knocking down all the isoforms of *SNRNP35* mRNA in HEK cells (Figure 3A-C). This study illustrated specificity of the *SNRNP35* knockdown, demonstrating a functional impact on U12 splicing of a target mRNA, *CHD1L* (Niemela et al., 2014), but not on U2 splicing of the same mRNA (Figure 3D-E).

**Figure 3.**
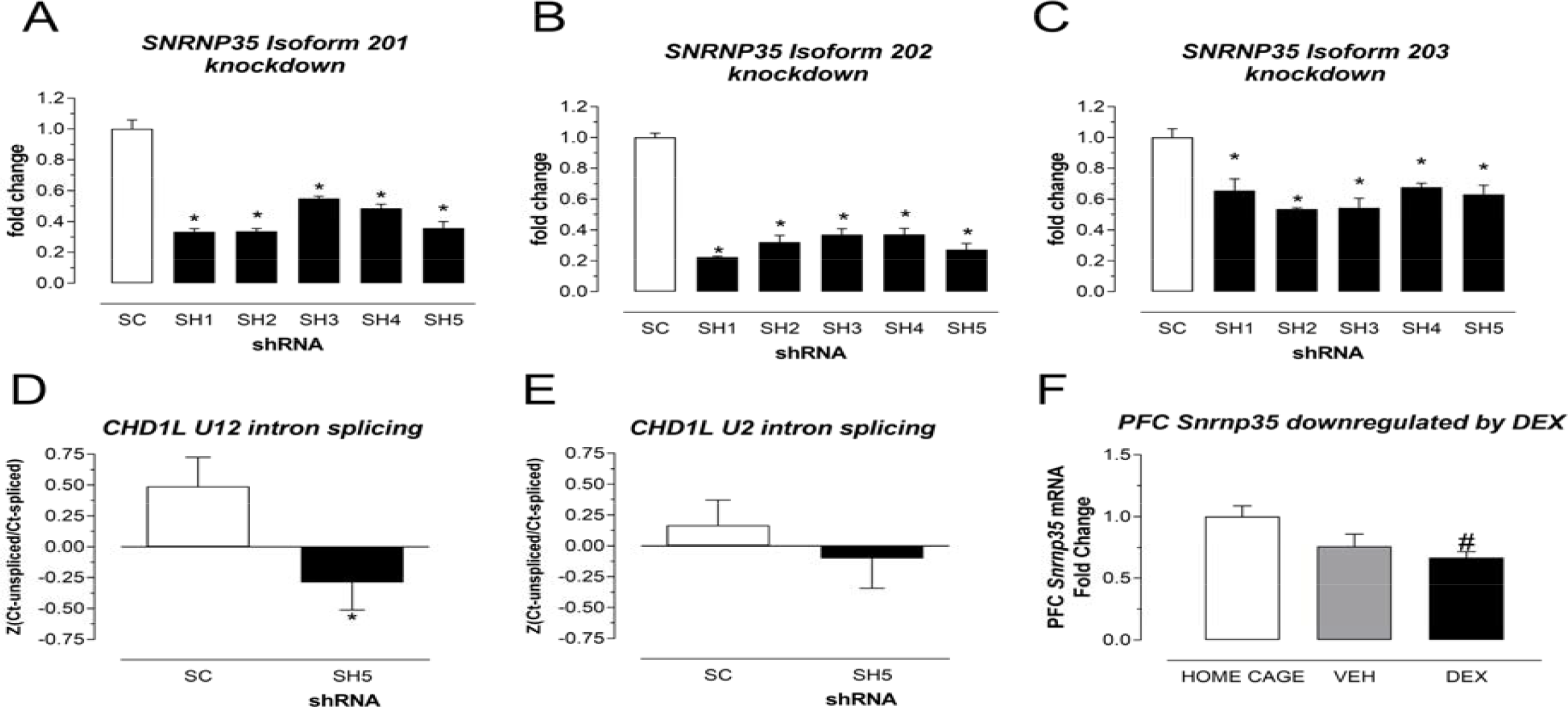
*SNRNP35* knockdown in human cells and mice. In HEK cells, the five short hairpin RNAs (shRNAs: SH1, SH2, SH3, SH4 and SH5) significantly down-regulated all the three protein coding *SNRNP35* RNA isoforms compared to scrambled (SC) RNA (A: for isoform 201 (hg38): p= 4.68×10^−05^, 4.71×10^−05^, 3.67×10^−04^, 2.36×10^−04^, and 1.29×10^−04^; B: for isoform 202 (hg38): p= 1.61×10^−07^, 1.20×10^−05^, 1.11×10^−05^, 1.23×10^−05^, and 5.47×10^−06^; C: for isoform 203 (hg38): p= 0.0106, 1.72×10^−04^, 1.61×10^−03^, 1.91×10^−03^ and 4.03×10^−03^). The SNRNP35 knockdown affected U12 (D), but not U2 splicing (E) of *CHD1L* target RNA. The repeated-measures ANOVA with technical replicate as within-subject factor and knock-down status as the between-subjects factor revealed an effect of knockdown status on U12 splicing F(1,17)=5.779; p=0.0279, and not on U2 splicing in F(1,15)=0.723, p=0.409). In mice dexamethasone (DEX) i.p. injection (10 mg/kg) downregulated prefrontal cortex (PFC) *Snrnp35* (Kruskal-Wallis H(2,21)=6.75, exact p= 0.0280; DEX vs. HOME CAGE adjusted p= 0.0303). *, vs. SC. #, vs. HOME CAGE.

### *SNRNP35* is downregulated in PFC by stress hormones

Given the effect of deployment stress on blood-based *SNRNP35* expression in the MRS study, we hypothesized that stress may also affect PFC *SNRNP35* expression. Stress hormones modulate gene expression through binding to the glucocorticoid receptor (GR), and subsequent binding to glucocorticoid-binding sequences (GBS). The mouse *Snrnp35* gene contains many GBS (15 sites out of a total of 196 transcription factor binding sites in the entire mouse gene; based on Gene Transcription Regulation Database (http://gtrd.biouml.org/) - **Table S13**). To examine the effect of stress-related GR activation on *SNRNP35* expression in a model system, we injected mice with 10 mg/kg dexamethasone (DEX), a synthetic stress hormone and potent GR-agonist. We observed significant *Snrnp35* downregulation in the PFC 4 hours later (DEX (n=7 males) vs. home cage (n=6 males: adjusted *P*-value = 0.0303), confirming regulation of *Snrnp35* by stress-hormones (Figure 3F). The direction of this stress-dependent effect on *Snrnp35* expression is consistent with the lower levels of *SNRNP35* in PTSD cases identified in both our GReX analysis and the MRS study.

## DISCUSSION

TI is a machine learning approach that translates GWAS findings into tissue-specific GReX associations with traits, adding gene, tissue and directional resolution. Applying this method to the multi-cohort PGC-PTSD GWAS, we have discovered two new putative PTSD susceptibility genes, *SNRNP35* and *SENP1*. Both genes are involved in post-transcriptional processes; *SNRNP35* is a subunit of the U11/12 minor spliceosome and involved in splicing of U12-type introns, and the SENP1 protein is a de-SUMOylation enzyme. *SNRNP35* mRNA is predicted to be downregulated in DLPFC in PTSD, a brain region of interest as it is involved in many stress-related neurobiological systems and processes (Averill et al., 2017; Nemeroff et al., 2006). Functional alterations in DLPFC have been described in PTSD, contributing to dysregulated circuit transmission and hypothalamus-pituitary-adrenal (HPA) axis function (Averill et al., 2017; Nemeroff et al., 2006). *SENP1* is predicted to be upregulated in the left heart muscle, fitting with substantial prevalence of cardiovascular disease in PTSD (Wolf and Schnurr, 2016) and the genetic overlap with cardiometabolic traits (Sumner et al.).

In our overall PTSD pathway analysis, we saw enrichments of only two small gene-sets (<20 genes). The first pathway is related to the activity of protein phosphatase 2A, which is an enzyme that has been shown to be increased in rat PFC and hippocampus after single or repeated immobilization stress (Morinobu et al., 2003). The second pathway is related to the levels of an antioxidant, uric acid; interestingly, antioxidant capacity has been recently studied in affective and anxiety disorders as a biomarker and treatment target (Black et al., 2018). The C-PTSD pathway analysis revealed an enrichment for genes derived from the PTSD literature. Furthermore, the enrichment of genes associated with circadian rhythms was expected given the wealth of PTSD research in this area, and the common presentation of PTSD with sleep dysfunction, including insomnia and nightmares. To name a few, studies revealed altered circadian rhythms of cortisol in individuals with PTSD (Yehuda et al., 1996), and association between PTSD and variants in circadian rhythm genes (Linnstaedt et al., 2018; Logue et al., 2013). Finally, the enrichment of the PSD-95 gene set aligns with substantial evidence supporting a role for PSD-95 in synapse-related dysfunction in several neuropsychiatric disorders (Penzes et al., 2011). The gene encoding PSD-95 is highly expressed in the brain (incl. pituitary; (GTEx Consortium, 2017)) and was also the top-gene in our overall PTSD meta-analysis.

A number of the M-PTSD specific functional enrichments are also highly plausible; for example, enrichment H3K27 acetylation peaks from a gene-set derived from DLPFC neurons (Girdhar et al., 2018), known to have a role in stress models and neuropsychiatric disorders (McEwen et al., 2015), including PTSD (Maddox et al., 2018). Our results highlight a potential shared genetic basis between olfaction and M-PTSD, in line with previous findings of differential olfactory identification in individuals with combat-related M-PTSD compared to healthy controls (Vasterling et al., 2000); olfactory triggers for PTSD intrusion symptoms (Daniels and Vermetten, 2016); olfactory-based treatments for PTSD (Aiken and Berry, 2015); and the key role of olfaction in fear conditioning in animal models (Morrison et al., 2015). Given the significant genetic overlap between olfaction and PTSD, our results support the hypothesis that differential sensitivity to odors may predispose to development of PTSD. Finally, the enrichment of gliogenesis, and glial and astrocyte differentiation in PTSD genetic susceptibility, is in concordance with the wealth of recent literature involving both these non-neuronal cell types with stress-related pathophysiology (Hodes et al., 2015; Sanacora and Banasr, 2013).

Both genes reaching study-wide significance (*SNRNP35, SENP1*) were significant in the M-PTSD, but not C-PTSD, analysis. Before interpreting M-PTSD and C-PTSD differences in genetic architecture, it is important to recognize a few methodological or statistical limitations. It is likely that the proxies used to delineate trauma type are imperfect, as inclusion of an individual in a military cohort does not preclude an experience of civilian trauma. These delineations also lacked nuance; for example, we do not distinguish between different types of military and civilian trauma or differences in the degree of trauma exposure. Civilian trauma is associated with more etiologic heterogeneity (clinical presentation differences among exposed individuals), while military trauma is related to more clinical heterogeneity (clinical presentation differences among those meeting PTSD diagnosis) (Prescott, 2012). Moreover, military cohorts may be more homogeneous in terms of gender (predominantly male), ancestry (mostly EA), and age than the civilian, and the lack of significance for these two genes in the transethnic or ancestry-specific C-PTSD analyses may be attributable to lack of power. We also note that there may be differences in control ascertainment between the two groups; whereas controls for military cohorts are trauma-exposed service members without PTSD, controls within civilian cohorts may be more diverse in the degrees of exposure trauma. Therefore, combining all civilian trauma studies may reduce the likelihood of identifying genes for C-PTSD-risk Finally, our C-PTSD analysis was based on raw genotype (Gamazon et al., 2015), while M-PTSD analysis was based on summary statistics (Barbeira et al., 2018). However, it is unlikely that these factors account entirely for the difference in effect sizes between C-PTSD and M-PTSD.

Sex differences in trauma exposure, symptom expression, levels of support, access to treatment, and treatment response may substantially affect PTSD outcomes (Breslau, 2002; Olff, 2017). The latest PGC-PTSD GWAS study has shown evidence for different SNP-heritability between men and women (Duncan et al., 2018). In the present study, we were able to investigate the contributions of sex to the genetic associations with PTSD in the military and civilian cohorts. Our comparisons of male-specific and female-specific C-PTSD and M-PTSD signals did not indicate any higher similarity between male-specific M-PTSD and male-specific C-PTSD compared to the similarity between male-specific M-PTSD and female-specific M-PTSD or male-specific C-PTSD and female-specific C-PTSD. Thus, the M-PTSD findings appear to be at least partially trauma type specific, rather than driven by sex. We urge that these questions be addressed in future large-scale PTSD GWAS; our analysis is relatively small, particularly with regards to military females, and we did not have access to raw genotype data for the military cohorts, precluding a more nuanced sex-specific analysis.

The M-PTSD and C-PTSD differences are intriguing and suggest that further research is necessary to elucidate the potentially different genetic etiologies of PTSD related to civilian and military trauma. To date, C-PTSD and M-PTSD have been considered the same disease, with comparable biological underpinnings. However, C-PTSD and M-PTSD sometimes differ in terms of clinical presentation, prevalence and environmental risk factors; for example, trauma severity, gender, age, race all have significantly different effect sizes between C-PTSD and M-PTSD. Little is known about the possible biological and molecular differences between C-PTSD and M-PTSD. Not surprisingly, biomarkers discovered in civilian studies are not always replicated in military studies (D. Norrholm and Jovanovic, 2011). For instance, genetic risk factors discovered in the largest PTSD-GWAS meta-analysis, which consisted of 82% civilian trauma samples, were not replicated in a large military sample, and vice versa (Duncan et al., 2018; Stein et al., 2016). Based on our results and previous findings, we hypothesize that C-PTSD and M-PTSD may have both shared and distinct genetic etiology, in line with subtypes of other complex, heterogeneous psychiatric disorders (Charney et al., 2017), or pairs of psychiatric disorders with substantial etiologic, symptomatic and diagnostic overlap (Bipolar Disorder and Schizophrenia Working Group of the Psychiatric Genomics Consortium, 2018). For these types of studies, substantial progress has been made by explicitly comparing cases of each disorder or subtype. However, given the limitations of our design, further work is needed to consolidate this hypothesis.

Although the gene-by-tissue associations identified point to biologically plausible tissues, we caution against over-interpretation of tissue sources in our study. The power of each tissue-specific predictor model (i.e., the accuracy of GReX prediction) is driven partially by the sample size of the expression quantitative trait loci (eQTL) reference panel and gene expression heritability in that tissue (Veturi and Ritchie, 2018). Additionally, there is a significant amount of eQTL sharing between similar tissues (e.g., within brain) (GTEx Consortium, 2017). It is also true that these predictor models are derived from bulk tissue and thus, tissue-specific weighs of SNPs might be driven by variations in cell type proportions between tissue-types or other tissue-specific factors. Importantly, using blood gene expression obtained by MRS (Breen et al., 2015), we observed concordance of predicted PTSD differences and observed PTSD differences with strong evidence for tissue-specificity (i.e. compared to the other tissues, our blood predictions matched better the observed blood differences).

Unlike traditional gene expression studies, TI approaches study only GReX. Any case-control differences identified are therefore due only to difference in allele frequencies, rather than influenced by environmental and/or epigenetic factors. These differences therefore cannot stem from differential exposure to trauma, or to any other environmental factors, or factors related to disease state - for example, from manifestation of symptoms or psychiatric medications. As TI models are derived from post-mortem adult tissues, the genotype-gene expression relationships encoded by these models will be biased by unknown stressors and other environmental factors in the lives of donors. As far as possible, we and others have controlled for these factors, including for example, correction for known diagnoses, age, post-mortem interval, smoking status, and surrogate variables (Gamazon et al.; Huckins et al., 2017), when constructing TI models. Although these methods may incompletely control for certain stressors, this will not lead to a systematic bias between cases and controls in our study, and as such should not lead to any inflation in our results.

Finally, we find significant evidence for downregulation of *SNRNP35* following deployment stress, primarily in PTSD cases. Thus, *SNRNP35* downregulation in not only associated with PTSD genetic susceptibility, but also with deployment stress, warranting further exploration. We first knocked-down *SNRNP35* gene expression in a cell-culture system to demonstrate that downregulation of this specific subunit of the minor spliceosome is sufficient to cause functional changes in the levels of U12 splicing. Since U12 splicing is not wide-spread in the genome (Turunen et al., 2013), *SNRNP35* downregulation is expected to have a finite number of directly affected downstream pathways that need to be tracked in post-mortem brains from trauma-exposed subjects with or without PTSD.

We further confirmed that the administration of high dose of a synthetic stress hormone, mimicking the glucocorticoid elevations after activation of the HPA-axis, can downregulate this gene in the mouse PFC. We used DEX, a potent GR agonist, and the observed *Snrnp35* downregulation is likely mediated though GR-binding at specific GBS of the gene. Previous studies have shown that DEX-administration, only at the high dose we used, can increase anxiety in the elevated-plus-maze, and that this elevation can be blocked by an opioid agonist (Vafaei et al., 2008) or can potentiate the hypermotility caused by opioids (Capasso et al., 1992). These observations open new avenues for future translational studies.

In conclusion, our analyses of GReX in PTSD identified novel genes for PTSD-risk, with a tissue resolution specific to military vs. civilian trauma. We identify *SNRNP35* as the most promising gene for further functional investigation of its trauma type specific role in vulnerability to and resilience against PTSD.

## ACKNOWLEDGMENTS

We thank study participants and research groups contributing to PGC-PTSD (Adam X Maihofer, Alicia R Martin, Allison E Ashley-Koch, Ananda B Amstadter, Andrea L Roberts, Anthony King, Bekh Bradley, Caroline M Nievergelt, Chia-Yen Chen, Daniel J Stein, Dewleen G Baker, Eric O Johnson, Henry R Kranzler, Hongyu Zhao, Israel Liberzon, Jean C Beckham, Jennifer A Sumner, Joel E Gelernter, John P Rice, Jonathan Bisson, Jordan W Smoller, Karestan C Koenen, Kerry J Ressler, Laramie E Duncan, Laura J Bierut, Lindsay A Farrer, Lynn M Almli, Mark J Daly, Mark W Logue, Mark W Miller, Melanie E Garrett, Michael A Hauser, Murray B Stein, Nancy L Saccone, Nastassja Koen, Nathan A Kimbrel, Nicole R Nugent, Rachel Yehuda, Rajendra A Morey, Robert J Ursano, Ronald C Kessler, Shareefa Dalvie, Stephan Ripke) for sharing their data.

### Funding

The PGC-PTSD data freeze related to this manuscript was supported by One Mind, Cohen Veterans Bioscience and the Stanley Center for Psychiatric Research. Harvard Medical School KL2/Catalyst Medical Research Investigator Training award, Brain & Behavior Research Foundation NARSAD Young Investigator Grant, and McLean Hospital Jonathan Edward Brooking Mental Health Research Fellowship (to NPD). U01 MH109536 (EAS and PS), 1R01MH118278-01 and Seaver Foundation Faculty Scholar Award (to LMH). R01MH107666, R01 MH101820, and P30 DK20595 (to HI). MRS and MRS-II (including RNA-seq) were funded by the Marine Corps, Navy Bureau of Medicine and Surgery (BUMED), VA Health Research and Development (HSR&D) and Navy BUMED.

### Data and materials availability

All data is available in the main text or the supplementary materials.

## AUTHOR CONTRIBUTIONS

Conceptualization (LMH, JDB, PS, EAS, NPD), data curation (LMH, MSB, CC, JH, TK, CK, MWL, AL, FGM, YP, LGS, SJv, DGB, LED, SJG, AXM, TJ, CMN, NPD), formal analysis (LMH, CC, NPD), funding acquisition (LMH, IL, KCK, CMN, JDB, PS, KJR, EAS, NPD), investigation (LMH, MSB, CC, JH, TK, ACd, AD, KG, GEH, CK, MWL, AL, FGM, HTN, YP, SJv, TJ, MK, MWM, SB, CMN, PS, KJR, EAS, NPD), methodology (LMH, MSB, CC, JH, TK, ACd, AD, KG, GEH, CK, AL, FGM, YP, SJv, NC, HI, KJR, EAS, NPD), project administration (LMH, NPD), resources (DGB, MAG, SJG, HI, VBR, MWM, CMN, PS, KJR, EAS, NPD), software (LMH, MSB, CC, AD, YP, HI, EAS, NPD), supervision (LMH, TK, TJ, MK, MWM, SB, CMN, JDB, PS, KJR, EAS, NPD), validation (LMH, MSB, CC, JH, TK, ACd, CK, MWL, AL, FGM, YP, SJv, TJ, CMN, KJR, NPD), visualization (LMH, MSB, CC, JH, ACd, NPD), writing – original draft (LMH, NPD), and writing – review & editing (LMH, MSB, CC, JH, TK, ACd, AD, KG, GEH, CK, MWL, AL, FGM, HTN, YP, DR, LGS, SJv, DGB, C-YC, NC, LED, MAG, SJG, HI, AXM, VBR, JWS, DJS, RY, IL, KCK, TJ, MK, MWM, SB, CMN, JDB, PS, KJR, EAS, NPD).

## METHODS

### Genotype data

Genotype data was obtained through the PTSD Workgroup of the Psychiatric Genomics Consortium (PGC-PTSD) data; https://pgc-ptsd.com/). Details regarding participants, genotyping, quality control, imputation, and ancestry assignment were reported previously (Duncan et al.). PGC-PTSD data used in this manuscript included 32,898 individuals (9,087 cases and 23,811 controls), partitioned according to self-defined ethnicity (broadly, in order of cohort size, European/European-American; African-American; Hispanic-American/Latin-American; South-African), and trauma type (military/civilian). Due to restrictions on sharing of raw data for some of these cohorts, our study includes analyses of both raw data, and summary statistics (**Table S1A**):

- Raw genotype data were available from seven civilian cohorts totaling 12,939 individuals (2,978 cases - 9,961 controls): Collaborative Genetic Study of Nicotine Dependence (COG) (Bierut et al., 2007) Family Study of Cocaine Dependence (FSCD) (Bierut et al., 2008); Yale-Penn, previously Genetics of Substance Dependence Study, (YP); Grady Trauma Project (GTP) (Kilaru et al., 2016); Nurses’ Health Study (NHS) (Koenen et al., 2009); Drakenstein Child Health Studies, two South African samples (SA) (Koen et al., 2014; Stein et al., 2015)
- Ancestry-specific summary statistics data were available from seven military cohorts prepared with a pipeline described in (Duncan et al., 2018). The total was 19,959 individuals (6,109 cases - 13,850 controls): Ohio National Guard (ONG) (Liberzon et al., 2014); Marine Resiliency Study (MRS) (Nievergelt et al., 2015); Mid-Atlantic Mental Illness Research Education and Clinical Center the study of Post-Deployment Mental Health Study (MIR6) (Liu et al., 2013); VA Boston-National Center for PTSD Study (NCPTSD) (Logue et al., 2013); two New Soldier Studies (NSS) within Army Study to Assess Risk and Resilience in Servicemembers (STARRS) (Stein et al., 2016); Pre/Post Deployment Study (PPDS) within Army STARRS (Stein et al., 2016).

For sex-specific analysis of the same civilian and military cohorts, we also had access to sex-specific summary statistics prepared with a pipeline described in (Nievergelt et al., 2018b). The breakdown of the sex-specific sample sizes can be found in **Table S1B**.

### Transcriptomic Imputation (TI)

We predicted genetically regulated gene expression (GReX) using publicly available predictor models derived from Genotype-Tissue Expression (GTEx) project (v6 release) and CommonMind Consortium (CMC) (v1 release) eQTL reference panels (Gamazon et al., 2015; Huckins et al., 2017). Briefly, predictor models were created from large, matched collections of genotype and gene expression data. Elastic net regression was used to identify SNPs within the cis-region (1Mb) that jointly predict the expression of a given gene. For each gene, dosages of SNPs included in the predictor model are weighted and combined to produce an estimate of genetically regulated gene expression. These predictor models may then be applied to genotype data, for example from GWAS studies.

For the PGC-PTSD cohorts for which we had access to raw data, we imputed genetically regulated gene expression (GReX) for each individual, across 11 brain regions, 5 cardiovascular tissues, 2 endocrine tissues, 1 peripheral nerve, 1 adipose tissue and whole blood, driven by prior hypotheses about the involvement of those tissues in PTSD (Averill et al., 2017; Daskalakis et al., 2018a; Nemeroff et al., 2006; Sareen, 2014) using PrediXcan (Gamazon et al., 2015). We then calculated associations between GReX and case-control status, correcting for ten genotype derived principal-components. For the PGC-PTSD, cohorts for which we only had access to summary statistics, we used S-PrediXcan (Barbeira et al., 2018). In this algorithm, summary statistics for SNPs within each gene model are combined, using model weights and LD between variants, to obtain genic association statistics. We have previously shown that the statistical calculations underlying PrediXcan (raw genotype) and S-PrediXcan (summary statistic based) are analogous (Barbeira et al., 2018; Huckins et al., 2017). Results from S-PrediXcan and PrediXcan are highly correlated (r≈.99) when applied to European populations, and high (r≈.92) in African American populations (Barbeira et al., 2018). Since we do not have any validation data for HA/LA ethnicities, and these military cohorts are relatively small (333 cases and 1,690 controls), we did not include genes reaching significance only in the HA/LA cohorts in our results.

### Meta-Analysis

We performed meta-analyses using an inverse variance based approach in METAL (Willer et al., 2010), for analyses (1) delineated by trauma type (Military/Civilian); and (2) delineated by ancestry. We note that, in this analysis, results from S-prediXcan are scaled to unit variance of a gene, whereas PrediXcan results are not. Therefore, an approach which combines PrediXcan and S-PrediXcan using inverse variance is inappropriate. For the ‘overall’ analysis we therefore combined cohorts using a sample size-based approach in METAL. We calculated “effective sample size”, N_eff_, according to the following formula (Willer et al., 2010): N_eff_ =4/(1/N_cases_+1/N_controls_). **Figure S1** illustrates our meta-analysis strategy.

We required that each meta-analysis included at least 1,000 cases. We applied two multiple-test corrections to ascertain significance, following previous PrediXcan literature (Barbeira et al., 2018; Huckins et al., 2017). First, a study-wide threshold, using a Bonferroni correction for all genes and tissues tested (p=2.36×10^−7^; based on 211,466 tissue-gene pairs, across all tissues). Second, a within-tissue significance threshold, accounting for all genes tested in each tissue (**Table S2**). It is likely that the study-wide threshold is overly conservative, given the high degree of eQTL sharing and gene expression correlation between genes and across tissues. Consequently, it is likely that we are performing far fewer independent tests than assumed under a Bonferroni correction. However, applying a less stringent threshold would risk identification of many false positive results, which we are careful to avoid.

### MRS validation of whole-blood results

The Marine Resiliency Study (MRS) is a prospective and longitudinal U.S. Marine cohort. The research team conducted structured clinical interviews on U.S. Marines and collected peripheral blood samples at 1-month prior-to deployment and 3-months following deployment to conflict zones (i.e. post-deployment). Details regarding the collection of clinical measures and peripheral blood samples have been described in detail previously^1-3^. Briefly, at the time of each blood draw, PTSD symptoms were assessed using a structured diagnostic interview, the Clinician Administered PTSD Scale (CAPS) and the PTSD Checklist (PCL). Diagnosis for PTSD was defined as a threat to life, injury, or physical integrity (Criterion A1) and the presence of at least one re-experiencing symptom and either three avoidance symptoms or two hyperarousal symptoms, or two avoidance symptoms plus two hyperarousal symptoms. Symptoms must have occurred at least once within the past month (frequency ≥ 1) and caused a moderate amount of distress (intensity ≥ 2).

All participants had to be symptom free with no PTSD diagnosis and a CAPS ≤ 25 at pre-deployment to be passed into subsequent gene expression analyses. Participants who fulfilled criteria for PTSD diagnosis were designated the PTSD group at post-deployment. Carefully matched trauma-exposed control samples with post-deployment CAPS ≤ 25 and those with matched post-deployment measures of combat exposure, age and ethnicity were designated the ‘trauma-exposed control’ group at post-deployment. Subsequently, if a Marine participant developed PTSD following trauma-exposure at 3-months post-deployment, their pre-deployment sample would be included in the ‘PTSD-risk’ group. Likewise, if a participant avoided PTSD symptoms at 3 months post-deployment their sample at pre-deployment was included in the ‘control’ group.

MRS gene expression data acquisition: Peripheral blood sample acquisition has been described in detail elsewhere (Breen et al., 2015; Glatt et al., 2013; Tylee et al., 2015). In brief, peripheral blood was obtained from U.S. Marine participants who served a seven-month deployment. Blood was drawn 1-month prior to deployment and again at 3-months post-deployment for each participant. Each blood sample (10ml) was collected into an EDTA-coated collection tube, RNA was isolated from peripheral blood leukocytes using LeukoLOCK Total RNA isolation and sequenced using the Illumina Hi-Seq 2000. From these samples, two separate data sets generated. The first data set of data included a total of 24 paired pre-deployment samples and 24 post-deployment samples, which were subjected to the Affymetrix Hu-Gene 1.0 ST Array. The second data set of data included a total of 130 pre-deployment samples and 134 post-deployment samples which were subjected to RNA-sequencing.

MRS gene expression data pre-processing: Data from each data set were processed, normalized and quality treated independently. Affymetrix arrays underwent robust multi-array average (RMA) normalization with additional GC-correction when possible [affy, oligo, gcrma (Wu et al., 2012)]. When multiple microarray probes mapped to the same HGNC symbol, the probes with the highest average expression across all samples was selected. RNA-sequencing were mapped and counted as described previously^1^. Genes with with >2 count per million (cpm) in at least 50% of all samples were retained and subsequently normalized using *VOOM* in *limma* (Ritchie et al., 2015), a variance-stabilization transformation method resulting in a normally distributed data matrix. For each data set, normalized data were inspected for outlying samples using unsupervised hierarchical clustering of samples (based on Pearson’s coefficient and average distance metric) and principal component analysis to identify potential outliers outside two standard deviations from these grand averages; ten outliers were removed in total. A total of 11,090 genes were expressed in both microarray and RNA-sequencing data sets, for which 6,295 genes (56.7%) also had predicted GReX expression. Combat batch correction (Leek et al., 2012) was applied to combine the two datasets and reduce systematic sources of variability other than case/control status, such as technical variability, forming the bases for subsequent case-control analytic comparisons.

MRS differential gene expression (DGE): DGE analysis was performed using the *limma* package (Ritchie et al., 2015) to detect relationships between diagnostic status and gene expression levels. The covariates ancestry (genetic PC1), age, traumatic brain injury (TBI), alcohol and nicotine were included in all models to adjust for their potential confounding influence on gene expression between main group effects.

### Gene-set enrichment tests

We performed gene-set enrichment tests using MAGMA (de Leeuw et al., 2015). We created a set of associations statistics using results from the overall, C-PTSD, and M-PTSD transethnic meta-analyses. For each set of results, we selected the best (most significant) *P*-value per gene, applying a Bonferroni correction to account for the number of tissues tested. We performed three gene-set enrichment analyses for each of our association statistics.

First, we tested for enrichment of genes from PTSD literature. We downloaded from PubMed (https://www.ncbi.nlm.nih.gov/pubmed/) the 700+ publication list according to “PTSD” & “gene” search (November 1, 2017). From these publications, we discarded the ones that were not original investigations. From the remaining 511 publications (**Table S5**), we were able to extract 143 unique gene symbols from the publication title irrespective if the reported findings were positive or negative. The most frequent being the serotonin transporter gene (SLC6A4 in 35 publications). Of these 143 genes, 103 were included in our transcriptomic imputation analyses. Second, we tested for enrichment of 92 hypothesis-driven pathways, including gene-sets associated with other psychiatric disorders, stress hormones, and genes with H3K4me3 or H3K27ac peaks in neurons, or non-neurons (Girdhar et al., 2018). Third, we tested 8,582 gene-sets collated from publicly available databases including KEGG, GO, REACTOME, PANTHER, BIOCARTA, and MGI. For all gene-set analyses, we included only gene-sets with at least ten genes and used the “competitive” *P*-value from MAGMA. We applied an FDR-correction within each experiment to correct for multiple testing.

### Gene Ontology (GO)

GO analysis was performed using the GO module of Enrichr (http://amp.pharm.mssm.edu/Enrichr/)

### Cell Culture Study

HEK293 cells (ATCC CRL-1573) were maintained under standard conditions in DMEM supplemented with 10% FBS and 1% Antibiotic-Antimycotic (all ThermoFisher Scientific) at 37C and 5% CO_2_ (vol/vol). For cell culture experiments, cells were seeded in 24 well plates at 35,000 cells/well. Transfection was performed the next day using Lipofectamine 2000 Transfection Reagent (ThermoFisher Scientific), following the manufacturer’s protocol. Cells were harvested 24 hours post transfection using TrypLE Express (ThermoFisher Scientific).

shRNA Construction. A shRNA plasmid against hsaSNRNP35 was constructed as follows: We purchased plasmid pshRNA containing a U6 promoter and a multiple cloning site followed by a mCherry gene driven by the PGK promoter from VectorBuilder Inc (Santa Clara, CA). Target sequences for hsaSNRNP35 were derived from https://www.invivogen.com/sirnawizard/ using default settings. We designed custom 58nt oligos (Table 1) with AgeI/EcoRI restriction sites, annealed them to generate double stranded DNA fragments and ligated this fragment into the AgeI/EcoRI sites of pshRNA to generate pshSNRNP35. Similar, a scrambled control was constructed. Restriction digest and Sanger Sequencing confirmed the resulting plasmid. All plasmids will be deposited at addgene (addgene.com). RNA extraction and qPCR. Total RNA extraction, reverse transcription, and qPCR for cell culture and animal dexamethasone experiment was performed as follows: Total RNA was isolated and purified using the Quick-RNA Miniprep Kit (Zymo Research, Irvine, CA) according to the manufacturer’s protocol. RNA concentration was measured with The Qubit 2.0 Fluorometer (ThermoFisher Scientific). RNA was reverse transcribed with the SuperScript IV First-Strand Synthesis System (ThermoFisher Scientific, Waltham, MA,), using random hexamer primers provided within the kit. cDNA was amplified on an Applied Biosystems ViiA7 Real-Time PCR System with Power SYBR Green PCR Master Mix (ThermoFisher Scientific, Waltham, MA). GAPDH was used as control. Data were analyzed using the ΔΔCt method unless otherwise stated. Primer combinations are given in **Table S14**.

### Mouse study

The dexamethasone experiment was performed on adult (9 weeks old) C57BL/6J male mice obtained from The Jackson Laboratory. Mice were group-housed in a temperature-controlled vivarium, with ad libitum access to food and water. Animals were maintained on a 12-h light/dark cycle (lights on at 7:30 am), with experimental procedures being performed during the light cycle. Mice were administered dexamethasone (Sigma, St Louis, MO, USA, catalog no. D1159) intraperitoneally (i.p.) at a dose of 10 mg/kg dissolved in saline (DEX, n=7). The injection volume was 125µl/25g. Vehicle treated mice (VEH, n=8) were injected with the same amount of saline. Injections were performed between 8:00 and 8:30 am. The i.p. injection per se represents a moderate stressor that is able to induce a stress response. Therefore, an additional group of mice serving as baseline control did not receive any injection or handling prior sacrifice (home cage, n=6). 4 hours after the injection, all mice were sacrificed by decapitation following quick anesthesia by isoflurane. Brains were removed, snap-frozen in isopentane at −40 °C, and stored at - 80 °C until further processing. All procedures conformed to National Institutes of Health guidelines and were approved by McLean Hospital Institutional Animal Care and use Committee. Whole PFC tissue punches were performed (1.78 to 1.34 mm anterior of bregma; **Figure S3**) based on the Mouse Brain Atlas.

Total RNA was isolated and purified using the Quick-RNA mini kit (Zymo research, Irvine, CA, USA, catalog. no. R1054) according to the manufacturer’s protocol. RNA templates were reverse transcribed into cDNA with the Superscript IV kit (Thermo Scientific, Waltham, MA, USA, catalog no. 18091200) and random hexamer primers. cDNA was amplified on an Applied Biosystems ViiA7 Real-Time PCR System with POWRUP SYBR Green Master Mix (Thermo Scientific, Waltham, MA, USA, catalog no. 4368706). Specific primers and GAPDH housekeeping primers were as follows: Snrnp35 (fwd. 5′ CGGTGGAAACGGTTTTTCT 3′; rev. 5′ CGGTCATGTGGGTCTTCATC 3′), GAPDH (fwd. 5′ TATGACTCCACTCACGGCAA 3′; rev. 5′ ACATACTCAGCACCGGCCT 3′). Ct values were normalized using the established delta-delta Ct method (2^−∆∆Ct^) and normalized to GAPDH Cts.

## SUPPLEMENTARY INFORMATION

**Table S1 (separate file).** GWAS sample sizes for the main meta-analyses (A) and the sex-specific meta-analysis (B).

**Table S2.** Within-tissue significance thresholds and study-wide significance threshold.

| TI model | N of predicted genes | Within-tissue Significance thresholds |
| --- | --- | --- |
| GTEX Adipose Subcutaneous | 10861 | $4.60 \times 10^{-6}$ |
| GTEX Adrenal Gland | 9289 | $5.38 \times 10^{-6}$ |
| GTEX Artery Aorta | 10150 | $4.93 \times 10^{-6}$ |
| GTEX Artery Coronary | 8871 | $5.64 \times 10^{-6}$ |
| GTEX Artery Tibial | 10647 | $4.70 \times 10^{-6}$ |
| GTEX Brain Anterior Cingulate Cortex BA24 | 8731 | $5.73 \times 10^{-6}$ |
| GTEX Brain Caudate Basal Ganglia | 9145 | $5.47 \times 10^{-6}$ |
| GTEX Brain Cerebellar Hemisphere | 9451 | $5.29 \times 10^{-6}$ |
| GTEX Brain Cerebellum | 10002 | $5.00 \times 10^{-6}$ |
| GTEX Brain Cortex | 9162 | $5.46 \times 10^{-6}$ |
| GTEX Brain Frontal Cortex BA9 | 9031 | $5.54 \times 10^{-6}$ |
| GTEX Brain Hippocampus | 8535 | $5.86 \times 10^{-6}$ |
| GTEX Brain Hypothalamus | 8551 | $5.85 \times 10^{-6}$ |
| GTEX Brain Nucleus Accumbens Basal Ganglia | 8913 | $5.61 \times 10^{-6}$ |
| GTEX Brain Putamen Basal Ganglia | 8759 | $5.71 \times 10^{-6}$ |
| GTEX Heart Atrial Appendage | 9438 | $5.30 \times 10^{-6}$ |
| GTEX Heart Left Ventricle | 9655 | $5.18 \times 10^{-6}$ |
| GTEX Nerve Tibial | 11440 | $4.37 \times 10^{-6}$ |
| GTEX Pituitary | 9159 | $5.46 \times 10^{-6}$ |
| GTEX Thyroid | 11176 | $4.47 \times 10^{-6}$ |
| GTEX Whole Blood | 10208 | $4.90 \times 10^{-6}$ |
| CMC Dorsolateral prefrontal cortex | 10292 | $4.86 \times 10^{-6}$ |
| Total | 211466<br>tissue-gene<br>pairs | $2.36 \times 10^{-7}$<br>(study-wide<br>significance<br>threshold) |

**Table S3 (separate file).** Pairwise enrichments and correlations of sex-specific results.

**Table S4.** Sex-specific Study-wide and within-tissue significant GReX associations with PTSD.

| Trauma-type (proxy) | Sex | Ancestry | Cases | Controls | Total | Tissue | Gene name | z-score | P-value |
| --- | --- | --- | --- | --- | --- | --- | --- | --- | --- |
| ALL | males | ALL | 5333 | 18118 | 23451 | Artery Tibial | *SLC25A36* | -4.76 | 1.96E-06 |
| ALL | males | EA | 3512 | 11769 | 15281 | Brain Frontal Cortex BA9 | ***SNRNP35*** * | -5.48 | 4.20E-08 |
| ALL | males | AA | 1488 | 4659 | 6147 | Adrenal Gland | *MYBPC2* | 4.71 | 2.46E-06 |
| MILITARY | males | ALL | 4578 | 14770 | 19348 | Heart Atrial Appendage | *SENPI* | 4.67 | 2.94E-06 |
| MILITARY | males | EA | 3367 | 10754 | 14121 | Brain Frontal Cortex BA9 | ***SNRNP35*** * | -5.67 | 1.41E-08 |
| MILITARY | males | EA | 3367 | 10754 | 14121 | Heart Atrial Appendage | *SENPI* | 4.64 | 3.39E-06 |
| CIVILIAN | females | ALL | 2391 | 5246 | 7637 | Brain Putamen Basal Ganglia | *IFNE1* | -4.81 | 1.54E-06 |
| CIVILIAN | females | ALL | 2391 | 5246 | 7637 | Adipose Subcutaneous | *HIST1H2AJ* | -4.69 | 2.72E-06 |

**Table S5 (separate file).** Extracted PTSD genes from the literature based on publication title.

**Table S6.** MAGMA gene-set enrichment in our overall, C-PTSD and M-PTSD transethnic meta-analyses.

| Transethnic Analysis based on Trauma-type | Gene-set type | Gene-set name | N genes | competitive P-values | FDR |
| --- | --- | --- | --- | --- | --- |
| ALL | publicly available gene sets | protein phosphatase type 2A regulator activity | 17 | 2.26E-06 | 0.019 |
| ALL | publicly available gene sets | Uric acid levels | 19 | 9.94E-05 | 0.014 |
| CIVILIAN | literature derived genes | Table S5 | 103 | 1.38E-03 | N/A |
| CIVILIAN | hypothesis driven | all.pli.sets.out:HIGH | 2715 | 9.98E-05 | 0.009 |
| CIVILIAN | hypothesis driven | all.pli.sets.out:MID | 3276 | 3.97E-04 | 0.018 |
| CIVILIAN | hypothesis driven | Circadian entrainment | 94 | 2.39E-03 | 0.073 |
| CIVILIAN | hypothesis driven | AUT-LoF | 113 | 5.78E-03 | 0.096 |
| CIVILIAN | hypothesis driven | CLOCK-CONTROLLED WEAK | 400 | 6.29E-03 | 0.096 |
| CIVILIAN | hypothesis driven | PSD-95 (core) | 56 | 5.34E-03 | 0.096 |
| CIVILIAN | publicly available gene sets | abnormal head shape | 53 | 9.35E-06 | 0.048 |
| CIVILIAN | publicly available gene sets | abnormal embryonic tissue morphology | 638 | 1.23E-05 | 0.048 |
| CIVILIAN | publicly available gene sets | shortened head | 27 | 1.98E-05 | 0.048 |
| CIVILIAN | publicly available gene sets | Early Phase of HIV Life Cycle | 12 | 2.22E-05 | 0.048 |
| CIVILIAN | publicly available gene sets | collagen | 82 | 5.54E-05 | 0.095 |
| MILITARY | hypothesis driven | H3K27ac.merged.PFC.neuronal. only | 1479 | 6.41E-04 | 0.059 |
| MILITARY | publicly available gene sets | Olfactory Signaling Pathway | 311 | 9.12E-08 | 0.001 |
| MILITARY | publicly available gene sets | regulation of astrocyte differentiation | 22 | 1.51E-07 | 0.001 |
| MILITARY | publicly available gene sets | positive regulation of glial cell differentiation | 24 | 2.36E-07 | 0.001 |
| MILITARY | publicly available gene sets | KEGG OLFACTORY TRANSDUCTION | 337 | 5.55E-07 | 0.001 |
| MILITARY | publicly available gene sets | regulation of glial cell differentiation | 45 | 4.67E-06 | 0.008 |
| MILITARY | publicly available gene sets | abnormal facial motor nucleus morphology | 13 | 7.21E-06 | 0.010 |
| MILITARY | publicly available gene sets | positive regulation of gliogenesis | 34 | 1.32E-05 | 0.016 |
| MILITARY | publicly available gene sets | decreased motor neuron number | 34 | 1.94E-05 | 0.021 |
| MILITARY | publicly available gene sets | no suckling reflex | 14 | 2.82E-05 | 0.027 |
| MILITARY | publicly available gene sets | regulation of protein tyrosine kinase activity | 48 | 3.64E-05 | 0.030 |
| MILITARY | publicly available gene sets | mediator complex | 32 | 3.85E-05 | 0.030 |
| MILITARY | publicly available gene sets | short snout | 80 | 4.42E-05 | 0.030 |
| MILITARY | publicly available gene sets | regulation of gliogenesis | 61 | 5.14E-05 | 0.030 |
| MILITARY | publicly available gene sets | regulation of peptidyl-tyrosine phosphorylation | 156 | 5.15E-05 | 0.030 |
| MILITARY | publicly available gene sets | abnormal pons morphology | 30 | 5.23E-05 | 0.030 |
| MILITARY | publicly available gene sets | regulation of RNA stability | 39 | 6.95E-05 | 0.037 |
| MILITARY | publicly available gene sets | positive regulation of isotype switching | 13 | 7.25E-05 | 0.037 |
| MILITARY | publicly available gene sets | positive regulation of cell development | 158 | 7.78E-05 | 0.037 |
| MILITARY | publicly available gene sets | KEGG PORPHYRIN AND CHLOROPHYLL METABOLISM | 38 | 1.14E-04 | 0.049 |
| MILITARY | publicly available gene sets | regulation of mRNA stability | 35 | 1.15E-04 | 0.049 |
| MILITARY | publicly available gene sets | chromatin DNA binding | 27 | 1.60E-04 | 0.063 |
| MILITARY | publicly available gene sets | regulation of isotype switching | 17 | 1.69E-04 | 0.063 |
| MILITARY | publicly available gene sets | mating | 32 | 1.70E-04 | 0.063 |
| MILITARY | publicly available gene sets | cerebral cortex cell migration | 27 | 1.80E-04 | 0.064 |
| MILITARY | publicly available gene sets | positive regulation of lymphocyte apoptotic process | 12 | 2.17E-04 | 0.074 |
| MILITARY | publicly available gene sets | abnormal snout morphology | 108 | 2.43E-04 | 0.080 |
| MILITARY | publicly available gene sets | positive regulation of DNA recombination | 15 | 2.53E-04 | 0.080 |
| MILITARY | publicly available gene sets | pseudouridine synthase activity | 12 | 2.71E-04 | 0.083 |

**Table S7 (separate file).** Observed PTSD-effects on leukocyte gene expression at pre- and post-deployment.

**Table S8A (separate file).** Post-deployment PTSD-effects on genetically-regulated gene expression (GReX) across tissues for MRS subjects with pre-deployment samples.

**Table S8B (separate file).** Post-deployment PTSD-effects on genetically-regulated gene expression (GReX) across tissues for MRS subjects with post-deployment samples.

**Table S9 (separate file).** Correlations of genetically-regulated gene expression (GReX) and observed leukocyte expression correlations at pre- and post-deployment.

**Table S10 (separate file).** Longitudinal (pre-to post-deployemt) analysis leukocyte gene expression.

**Table S11 (separate file).** Gene-ontology terms for the *SNRNP35* containing Module in control subjects from DLPFC RNA-seq dataset (Fromer et al., 2016).

**Table S12.** Top-10 Gene-ontology terms for the *SNRNP35* containing Module in control subjects from DLPFC RNA-seq dataset (Fromer et al., 2016)

| Term | Overlap | P-value | Adj. P | Genes |
| --- | --- | --- | --- | --- |
| RNA binding (GO:0003723) | 46/1388 | 1.56 x10<sup>-18</sup> | 2.78 x10<sup>-16</sup> | *RBM8A; CCDC47; RPL11; RPL36A; HMGB3; PARK7; HTATSFI; THG1L; RPS4X; MRPL20; ZMAT2; SNRPD1; BMS1; TMSB4X; MAGOH; SAP18; SNIP1; DENR; SNRPD3; ZNF622; SNRNP35; NPM1; NCBP2; ALYREF; NGDN; RPL22; RPS6; PDAP1; APTX; HARS2; MRPS5; NAP1L4; SNW1; MFAP1; DKC1; SYF2; THRAP3; CSDE1; RPL27A; HNRNPH2; PIN4; HNRNPC; SRSF5; KPNB1; RPS24; EIF3B* |
| mRNA processing (GO:0006397) | 17/284 | 4.29x10<sup>-11</sup> | 2.01 x10<sup>-08</sup> | *PPIL1; RBM8A; NCBP2; ALYREF; HTATSFI; PRPF18; SNW1; ZMAT2; SNRPD1; SYF2; MAGOH; HNRNPH2; HNRNPC; SNRPD3; SRSF5; SNRNP35; CTNNBL1* |
| RNA splicing, via transesterification reactions with bulged adenosine as nucleophile (GO:0000377) | 16/237 | 2.76 x10<sup>-11</sup> | 2.01 x10<sup>-08</sup> | *PPIL1; RBM8A; NCBP2; ALYREF; HTATSFI; SNW1; ZMAT2; SNRPD1; SYF2; MAGOH; HNRNPH2; HNRNPC; SNRPD3; SRSF5; SNRNP35; CTNNBL1* |
| mRNA splicing, via spliceosome (GO:0000398) | 16/262 | 1.23 x10<sup>-10</sup> | 3.84 x10<sup>-08</sup> | *PPIL1; RBM8A; NCBP2; ALYREF; HTATSFI; SNW1; ZMAT2; SNRPD1; SYF2; MAGOH; HNRNPH2; HNRNPC; SNRPD3; SRSF5; SNRNP35; CTNNBL1* |
| gene expression (GO:0010467) | 18/412 | 1.84x10<sup>-09</sup> | 4.29 x10<sup>-07</sup> | *RBM8A; GTF3C6; NCBP2; ALYREF; RPL11; RPL22; RPS6; MRPS18B; RPL36A; HARS2; MRPS5; RPS4X; MRPL51; DKC1; RPL27A; MAGOH; SRSF5; RPS24* |
| nuclear-transcribed mRNA catabolic process (GO:0000956) | 12/175 | 7.74 x10<sup>-09</sup> | 1.45 x10<sup>-06</sup> | *RPS4X; RBM8A; NCBP2; THRAP3; CSDE1; RPL27A; MAGOH; RPL11; RPL22; RPS6; RPL36A; RPS24* |
| cytosolic part (GO:0044445) | 11/160 | 3.25 x10<sup>-08</sup> | 1.46 x10<sup>-06</sup> | *RPS4X; RPL36AL; RPL27A; PSMC2; RPL11; RPL22; RPS6; RPL36A; ZNF622; MRPS5; RPS24* |
| U2-type spliceosomal complex (GO:0005684) | 8/61 | 1.66 x10<sup>-08</sup> | 1.46 x10<sup>-06</sup> | *PPIL1; PRPF18; SNW1; SNRPD1; SYF2; SNRPD3; HTATSFI; SNRNP35* |
| cytosolic ribosome (GO:0022626) | 10/125 | 3.32 x10<sup>-08</sup> | 1.46 x10<sup>-06</sup> | *RPS4X; RPL36AL; RPL27A; RPL11; RPL22; RPS6; RPL36A; ZNF622; MRPS5; RPS24* |
| nuclear-transcribed mRNA catabolic process, nonsense-mediated decay (GO:0000184) | 10/113 | 1.25 x10<sup>-08</sup> | 1.95 x10<sup>-06</sup> | *RPS4X; RBM8A; NCBP2; RPL27A; MAGOH; RPL11; RPL22; RPS6; RPL36A; RPS24* |

**Table S13 (separate file).** Mouse Snrnp35 transcription factor binding sites according to Gene Transcription Regulation Database.

**Table S14.**
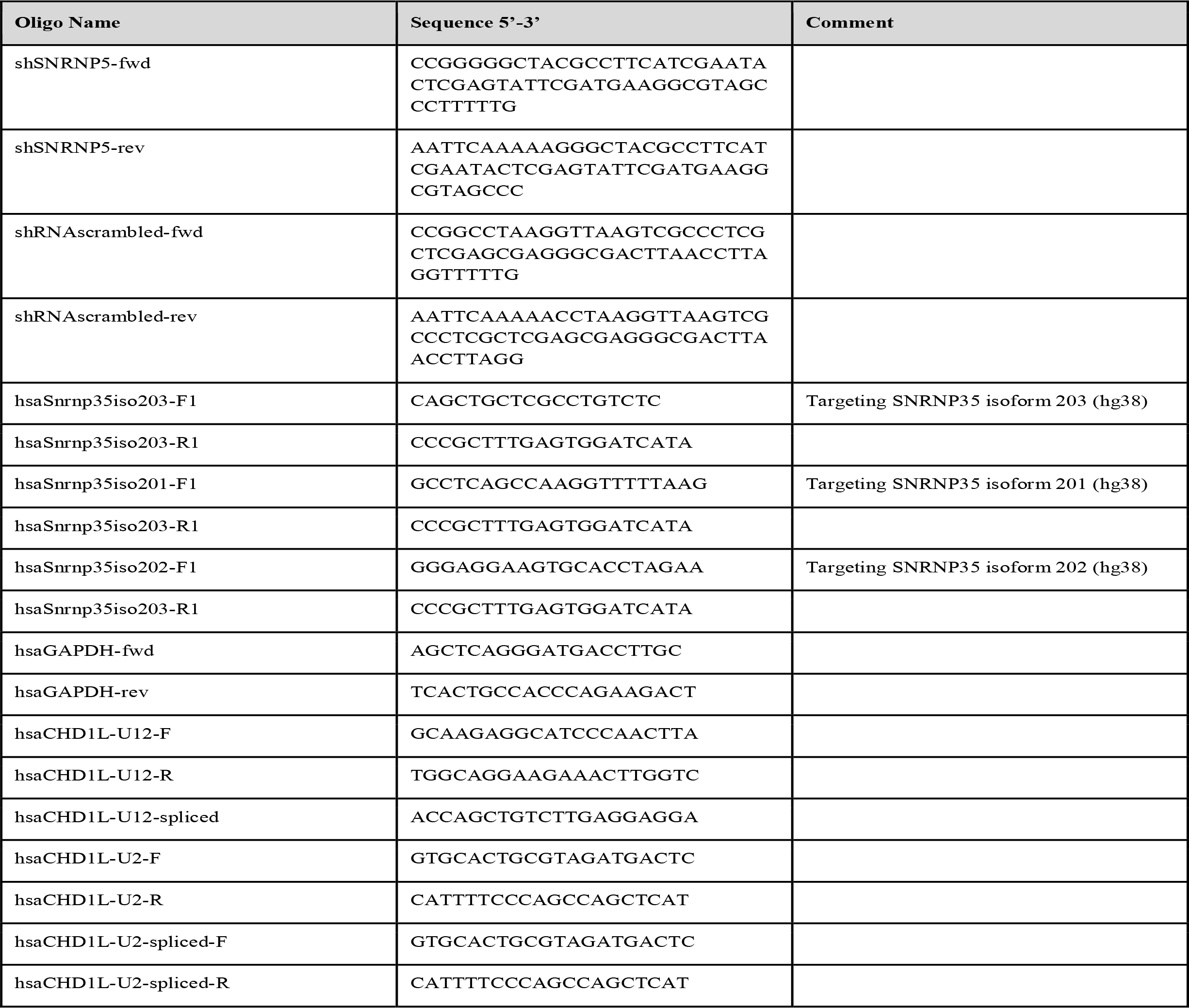
Primers used in cell culture.

**Figure S1.**
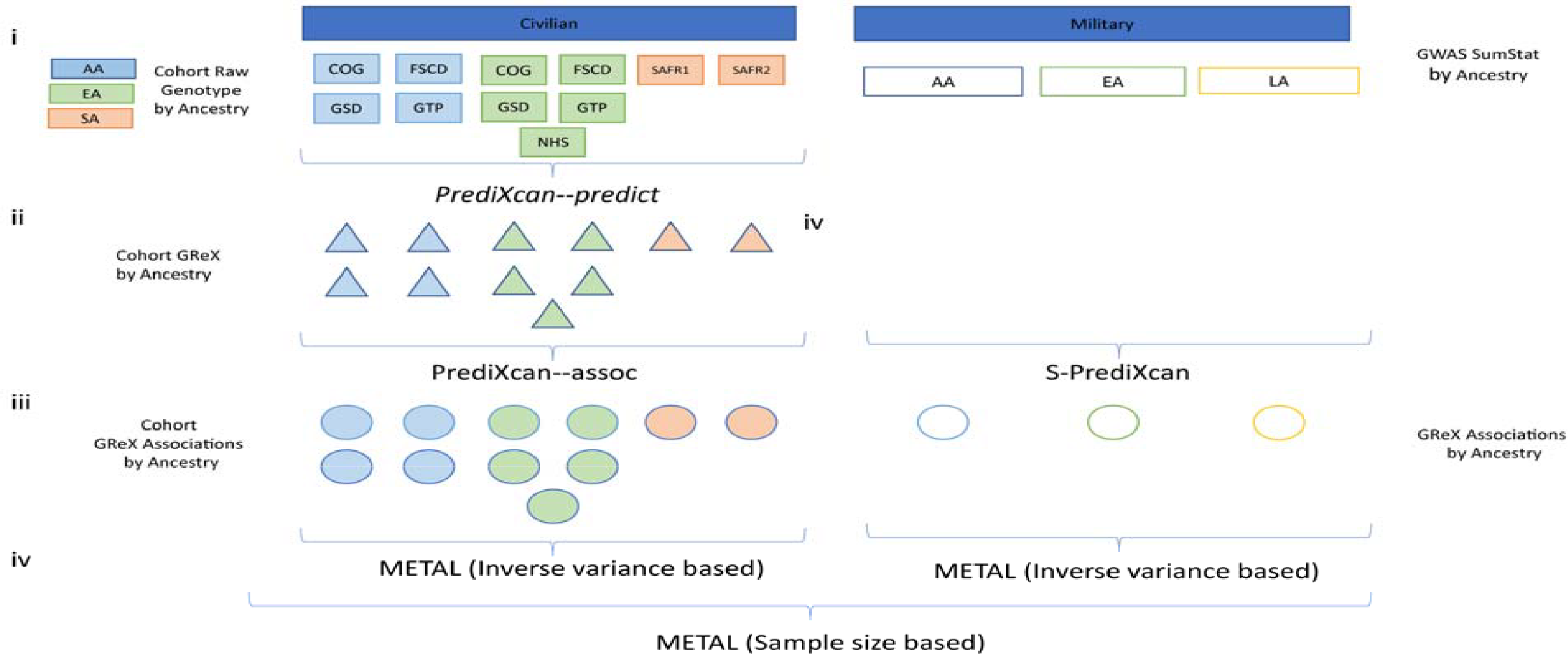
Analytical approach for calculation of Genetically regulated gene expression (GReX) associations statistics, and meta-analysis. (i) 11 ancestry-specific civilian cohorts with full raw genotype data (**Table S1**) were available (full-colored rectangles; blue: AA, green: EA and pink: SA). Data access restrictions meant that raw data were not available for military cohorts. Instead, we used three population-specific summary statistics (empty-colored rectangles: blue outline: AA, green outline: EA, and yellow outline: LA). (ii) Genetically regulated gene expression (GReX) was calculated across all 11 civilian cohorts with raw genotype data available (iii) GReX-PTSD associations were calculated for all 11 civilian cohorts (iv) GReX summary statistics were calculated for each set of Military ancestry-specific summary statistics, using S-PrediXcan (v) Within-trauma meta-analyses were performed using an inverse-variance based approach in METAL. Since effect sizes in S-PrediXcan were scaled to unit variance, civilian and military cohorts were meta-analyzed using a sample-size based approach.

**Figure S2.**
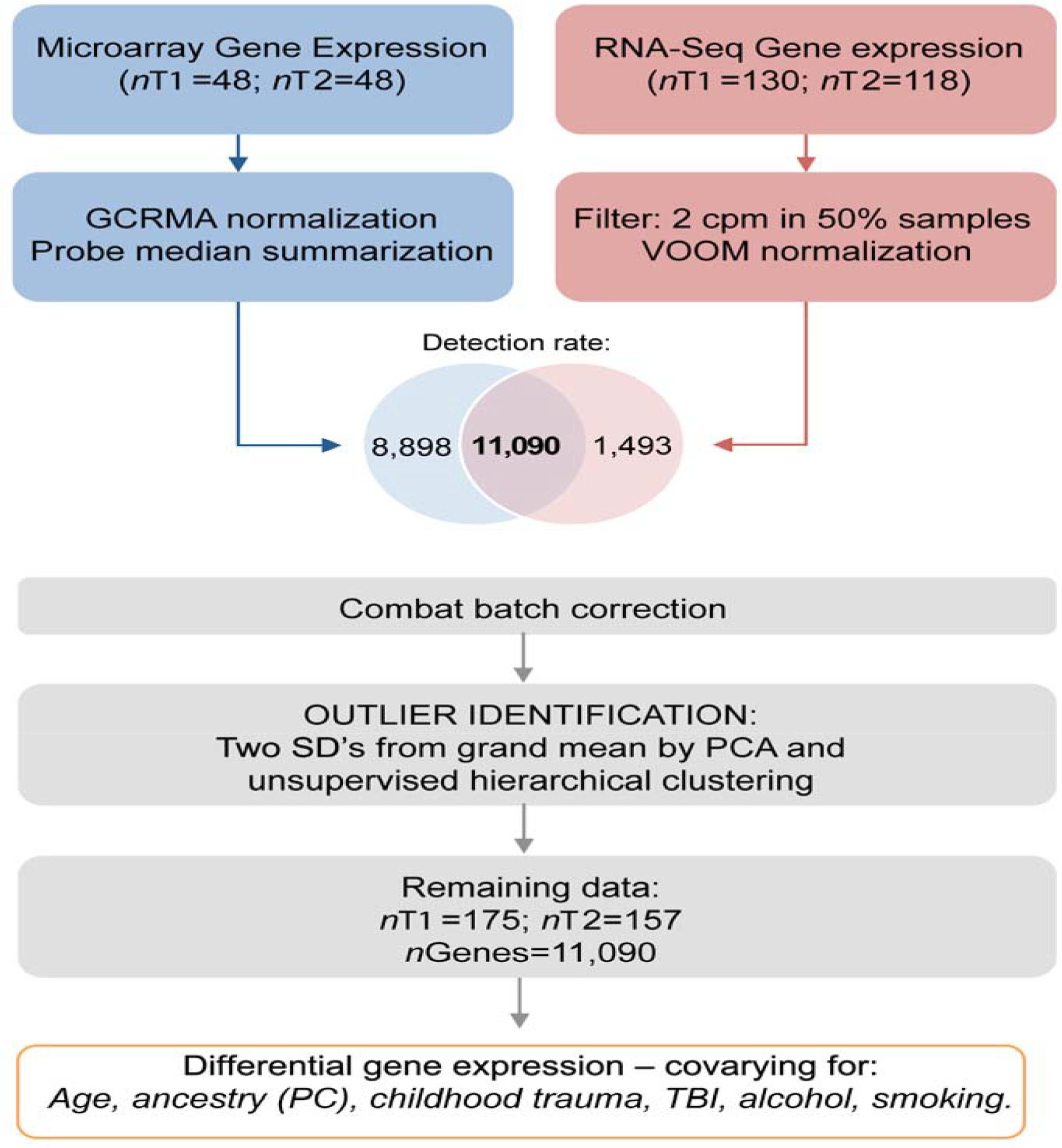
Analytical approach for quality control, batch correction, and differential gene expression in the Marine Resiliency Study.

**Figure S3.**
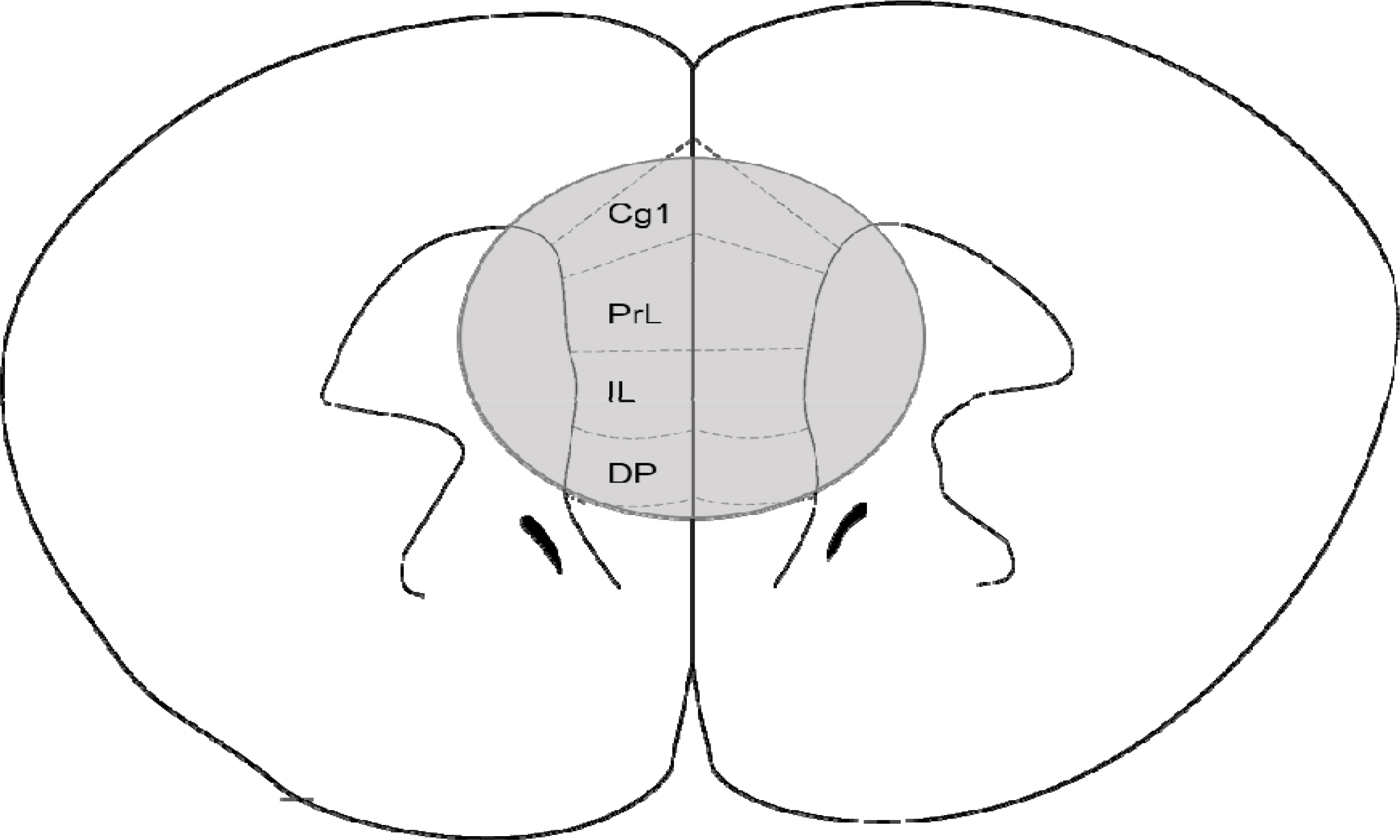
The area of the PFC mouse punch. Cg1: cingulate cortex, area 1; PrL: prelimbic cortex; IL: infralimbic cortex; DP: dorsal peduncular cortex.

